# MATAM: reconstruction of phylogenetic marker genes from short sequencing reads in metagenomes

**DOI:** 10.1101/141176

**Authors:** Pierre Pericard, Yoann Dufresne, Samuel Blanquart, Hélène Touzet

**Affiliations:** CRIStAL (UMR CNRS 9189, Université Lille 1), 59650 Villeneuve d’Ascq, France; Inria Lille Nord-Europe, 59650 Villeneuve d’Ascq, France

## Abstract

**Motivation:** Advances in the sequencing of uncultured environmental samples, dubbed metagenomics, raise a growing need for accurate taxonomic assignment. Accurate identification of organisms present within a community is essential to understanding even the most elementary ecosystems. However, current high-throughput sequencing technologies generate short reads which partially cover full-length marker genes and this poses difficult bioinformatic challenges for taxonomy identification at high resolution

**Results:** We designed MATAM, a software dedicated to the fast and accurate targeted assembly of short reads sequenced from a genomic marker of interest. The method implements a stepwise process based on construction and analysis of a read overlap graph. It is applied to the assembly of 16S rRNA markers and is validated on simulated, synthetic and genuine metagenomes. We show that MATAM outperforms other available methods in terms of low error rates and recovered genome fractions and is suitable to provide improved assemblies for precise taxonomic assignments.

**Availability:** https://github.com/bonsai-team/matam

## 1 Introduction

Shotgun metagenomic sequencing provides an unprecedented opportunity to study uncultured microbial samples, with multiple applications ranging from the human microbiome to soil or marine samples, for which the vast majority of microorganisms diversity remains unknown [13].

A major goal of metagenomic studies is to characterize the microbial diversity and ecological structure. This is often achieved by focusing on one of several phylogenetic marker genes [12, 28], that are ubiquitous in the taxonomic range of interest and exhibit variable discriminative regions. For bacterial communities, the gold standard marker is the 16S ribosomal RNA (rRNA, ~1500bp avg. length), for which millions of sequences are available in curated reference databases, such as Silva [24], RDP [2] or GreenGenes [3]. Traditionnal approaches such as amplicon sequencing are limited to the analysis of small portions of the marker sequences. This leads to strong technological limitations for organisms identification at sufficiently precise taxonomic levels, typically beyond genus [23]. To assign marker sequences to species, or even strains, we need to be able to recover full length rRNA with less than a few errors per kilobase. Metagenomic assemblers are not suitable for this task, because they are optimized to deal with whole genomes, and struggle to differentiate between very similar sequences [27]. To this respect, marker-oriented methods such as EMIRGE [19] and REAGO [33] were recently developed in order to assemble metagenomic read subsets into full length 16S rRNA contigs, thus aiming to improve the taxonomic assignment accuracy of environmental samples. EMIRGE uses a Bayesian approach to iteratively reconstruct 16S rRNA full length sequences. REAGO identifies rRNA reads using Infernal [20], and then constructs an overlap graph by searching for exact overlaps between reads using a suffix/prefix array. However, such tools still show some limitations in terms of recovery error rates as well as dealing with low abundance species.

In this work, we present MATAM, a new approach based on the construction and exploitation of an overlap graph, carefully designed to minimize the error rate and the risk of chimera formation. MATAM was validated on both simulated and actual sequencing data. It is able to reconstruct nearly full length 16S rRNAs and is robust to variations in the sequencing depth as well as community complexity.

## 2 Methods

### 2.1 Overview of MATAM

The MATAM (Mapping-Assisted Targeted-Assembly for Metagenomics) pipeline takes as input a set of shotgun metagenomics short reads and a reference database containing the largest possible set of sequences from a given target marker gene. MATAM identifies reads originating from that marker, and assembles nearly full length sequences of it. It is composed of four major steps illustrated in Figure 1. Although this method should work for any conserved and widely surveyed gene, we will focus on the 16S rRNA for the remainder of the article. Additional technical details and parameters are available in the Supplementary Methods.

**Figure 1:**
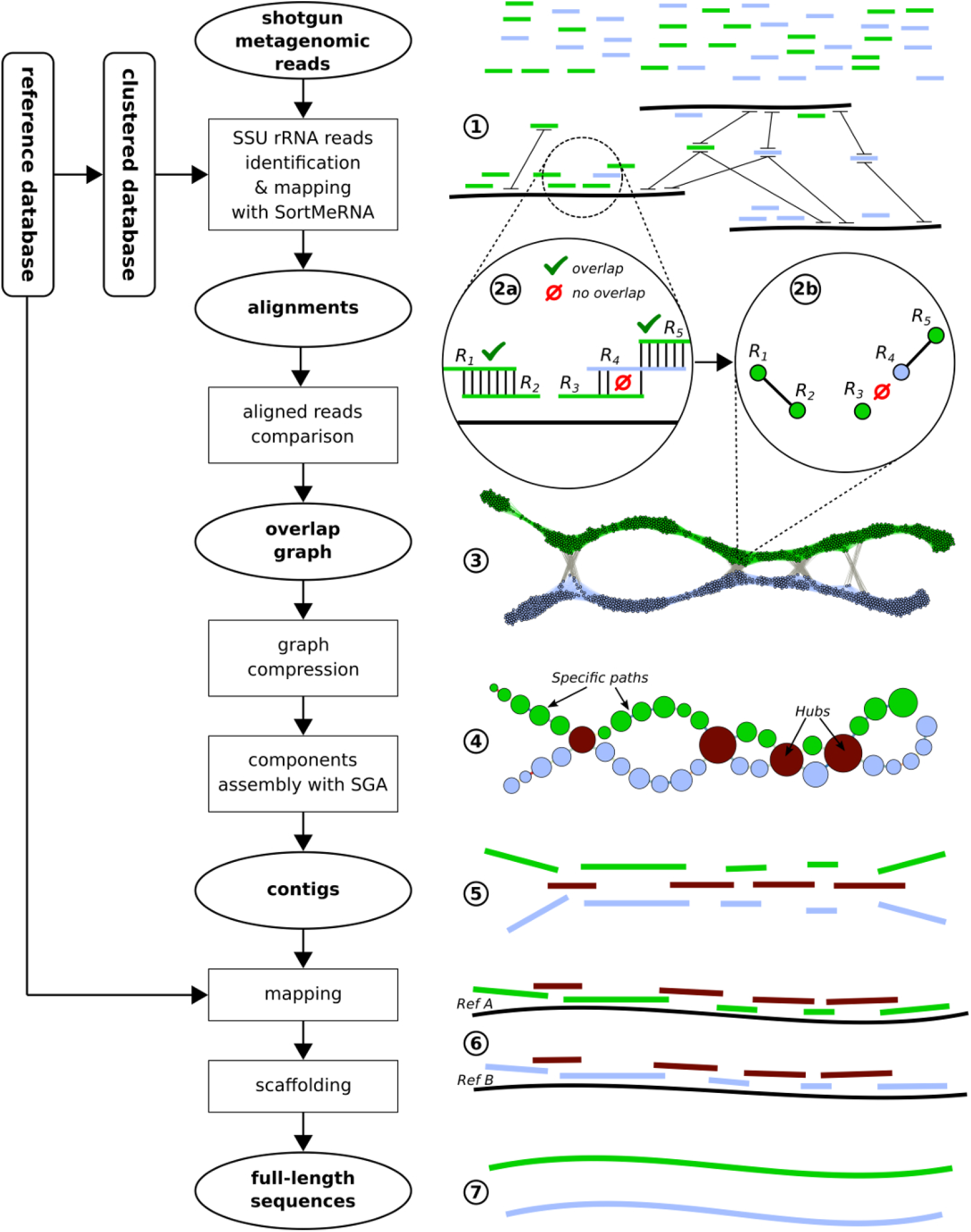
MATAM overview. On the left, we describe the main steps of the pipeline. On the right, we illustrate those steps when the sample contains two species (the first one in blue, the latter one in green). Starting from shotgun metagenomic reads, (1) we first identify SSU rRNA reads and align them on up to 10 sequences from a clustered reference database. (2a) Reads alignments are compared between them to compute reads pairwise alignments. (2b) An overlap with 100% identity between two reads corresponds to an edge in the (3) read overlap graph. (4) Using a breadth-first search, the overlap graph is then simplified into a compressed graph and subgraphs (hubs, specific paths, singletons) are identified. (5) Reads from each subgraph are assembled into contigs with SGA. (6) Contigs are aligned on the complete reference database and alignments are selected using a greedy algorithm. (7) Contigs aligned on the same reference are then scaffolded into full-length sequences.

### 2.2 Reference database construction

The availability of a reference database for the marker gene is an essential feature of the method, because it allows us to model the target sequences. For applications to 16S rRNA assembly, MATAM utilizes Silva 128 SSU Ref NR database [24]. From this reference database that we denote as *complete*, we also build a *clustered* reference database, that provides a coarse-grained representation of the taxonomic space. For that task, we use Sumaclust [17, 8] using a 95% identity threshold.

### 2.3 rRNA reads identification and mapping

In the first step, reads are mapped against the clustered reference database using SortMeRNA [10, 9]. This step allows to quickly sort out 16S rRNA reads from the whole set of reads, providing high quality alignments. For each read, we keep up to ten best alignments against the reference database. Moreover, this mapping step yields a broad classification of the 16S rRNA reads. Indeed, reads coming from distantly related species are aligned against their respective closest known references, which nest in distant lineages of the taxonomy, while reads from closely related species are aligned against closely related references.

### 2.4 Construction of the overlap graph

The identified 16S rRNA reads are then organized into an *overlap graph* defined as follows: graph nodes are reads, and an undirected edge connects two nodes if the two reads overlap with a sufficient length and with a sufficient identity to assert that they originated from a common sampled taxon. The standard approach to build such an overlap graph requires comparison of each read with each other, which is time-consuming. Here, we use alignment information to sort through candidate read pairs in a very efficient manner. For each pairing, we consider only reads that share alignments with at least one common reference sequence and for which the alignments are overlapping on more than 50 nucleotides with 100% identity. This strict criterion allows us to reduce the risk of connecting reads from unrelated taxa, which would in turn produce chimeras. By doing so, we discard reads containing sequencing errors in their overlap, which is bearable considering the nowadays very low sequencing error rates of short reads.

### 2.5 Extracting contigs from the overlap graph

Although the overlap graph appears very bushy, it also reveals some general trends. While it exhibits highly connected subgraphs, it also displays disjoint paths (see Figure 1 for an example). We simplify the graph by performing a breadth first traversal starting from a random node to annotate the nodes with their depth. All nodes with equal depth that are connected in a single connected component are collapsed into a single *compressed node* and outgoing edges are merged into a *compressed edge.* Low support compressed nodes containing a single read, and compressed edges representing a single overlap are removed. The resulting graph, called the *compressed graph*, is several order of magnitude smaller than the initial overlap graph. We partition this graph in three categories of subgraphs: *hubs*, that are nodes with an degree strictly greater than two, *specific paths* that are sequences of nodes of degree two or one, and *singletons* that are non-connected nodes. Intuitively, hubs correspond to the highly connected subgraphs in the overlap graph, and are likely to contain mainly reads coming from conserved regions shared in many species, thus overlaping without error even for distantly related taxa. Specific paths tend to contain reads originating from variable regions of the 16S gene, that are specific to one or few closely related species. For each subgraph in the compressed graph (hubs, specific paths, singletons), we extract the underlying sets of reads and build an individual assembly using the genomic assembler SGA [30]. Note that any other state-of-the art genomic assembler could be used here. As a result, we obtain one or more contigs for each subgraph.

### 2.6 Contigs scaffolding

We use a greedy algorithm to scaffold the contigs obtained in the previous step. For that task, contigs are first mapped against the complete reference database, and all alignments within the 1% range of suboptimal scores are kept. We then select contigs by increasing number of matches and decreasing lengths. By doing so, a long contig with a unique alignment will be selected for scaffolding before a short contig exhibiting a large number of alignments. Such long contig can be assigned non-ambiguously to a single species, while the short contig with multiple matches rather corresponds to a conserved region of the marker and is used to fill in the blanks between the specific contigs. Contigs matching against the same reference sequence are then merged into a single consensus scaffold. Redundant scaffolds included in larger ones are removed. Finally, only scaffolds larger than 500bp are retained. This yields the final MATAM output which could be used for the purpose of taxonomic assignment.

## 3 Implementation

MATAM was implemented in Python 3, except for the overlap graph building and compression steps that were written in C++11 using the SeqAn library [4], and is available *via* Docker and Conda. MATAM is distributed under the GNU Affero GPL v3.0 licence and the source code is freely available at the following URL: https://github.com/bonsai-team/matam. All MATAM runs presented in this article were performed using MATAM v0.9.9.

## 4 Results

MATAM performance was compared with those of two general-purpose metagenomic assemblers, SPAdes [1, 21] and MEGAHIT [11], as well as with two methods specialized in 16S rRNA assembly, EMIRGE [19] and REAGO [33]. The five tools were run on three different datasets, chosen for their complementarity and the possibility to validate the reconstructed candidate 16S rRNA sequences: a simulated dataset [15], a synthetic microbial community [29], and two environmental samples from human gut and mouth providing amplicon based taxonomic assignments [31]. SortMeRNA was used to extract 16S rRNA reads from these datasets before assembling them with SPAdes and MEGAHIT. Complete command-lines and parameters are available in the Supplementary Results.

In order to compare the five methods on a common ground, the same validation procedure was applied for all experiments. Only reconstructed sequences with lengths exceeding 500bp were considered, and chimeric sequences were filtered out by the UCHIME algorithm [5] implemented in VSEARCH [25] and querying the Silva 128 SSU Ref Nr99 database. For each experiment, we indicate the proportion of chimeric contigs (*% chimeras*, which is the total size of all chimeric contigs divided by the assembly total size). All the following measures were then computed on the remaining assemblies. When the sequences present in the sample are actually known (see Sections 4.1 and 4.2), the assembly quality assessment was performed with MetaQuast [18] by aligning the contigs against the original sample sequences, and considering the following metrics: the *number of contigs* (#contigs), which is the total number of contigs of lengths greater than 500bp; the *total length* (TL), which is the total number of bases in the contigs; the *total aligned length* (TAL), which is the total number of aligned nucleotides in the contigs; the *genome fraction* (GF), which stands for the total number of nucleotides from the original sample sequences covered with contigs divided by the total size of the sample sequences; the *error rate* (ER), which consists in the percentage of observed mismatches and indels with respect to the closest matched sequence in the original sample. Finaly, taxonomic assignments were carried out with the RDP Classifier [32]. The assemblies evaluation protocol, command-lines and parameters can be found in the Supplementary Results.

### 4.1 Simulated metagenomic datasets with varying sequencing depth

In the first experiment, we evaluated the ability of methods to correctly reconstruct the 16S rRNA sequences in the context of low sequencing depth. For that, we used a selection of 122 genomes providing a realistic taxomical diversity [15, 22], that contains 287 distinct 16S rRNA copies. We generated five datasets with varying sequencing depths: 50x, 20x, 10x, 5x and 2x per genome. Illumina reads were simulated with the ART simulator [7], using the HiSeq2500 built-in error profile, 101bp read length, and 250bp fragment length with a 30bp standard-deviation (SD). In this simulation, all species are equally distributed, which corresponds to the *high complexity community* introduced in [15]. Simulation command-line and parameters can be found in the Supplementary Material (section 4.3.1).

Table 1 shows the results averaged over the five datasets *(mean* metrics and their respective standard deviation, SD). More than 99% of the MATAM sequences were aligned by MetaQuast to one of the 287 16S rRNA sequences from the initial sample (mean TAL/TL), while among other methods, this proportion reached at best 91%, with REAGO. Congruently, MATAM sequences obtained the lowest average error rate (ER=0.03%), which represents more than a ten-fold accuracy gain compared to the other assemblers, and a twenty-fold improvement over EMIRGE. Furthermore, EMIRGE sequences contained 0.5% of unknown nucleotides (Ns), bringing its effective ER above 1%. Additionally, MATAM recovered about thirty times less chimeras than REAGO and EMIRGE did.

**Table 1:**
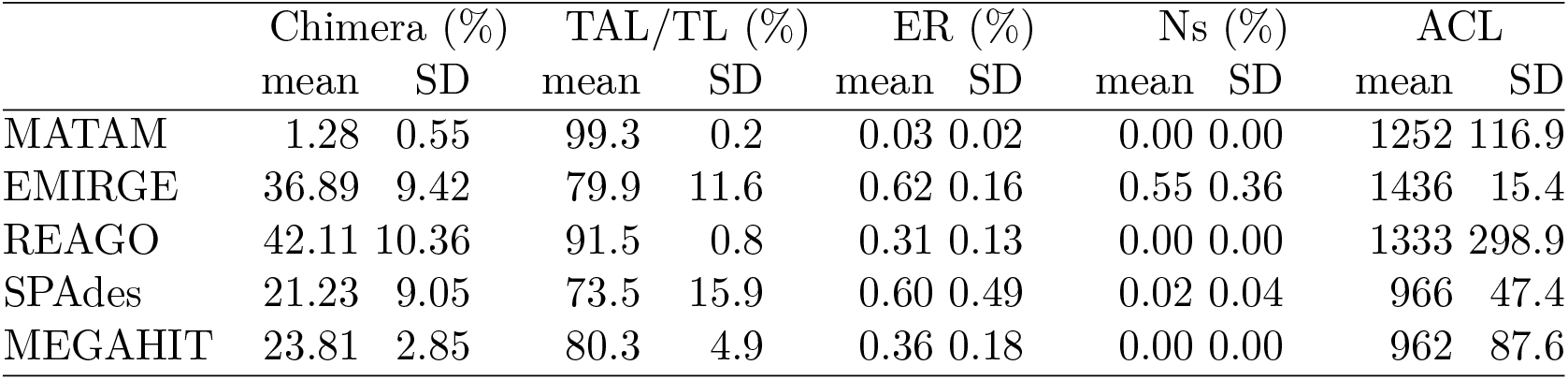
Results for the simulated dataset with varying sequencing depth. We provide averaged metrics for the five sequencing depths. ACL is the average contig length.

|  | Chimera (%) |  | TAL/TL (%) |  | ER (%) |  | Ns (%) |  | ACL |  |
| --- | --- | --- | --- | --- | --- | --- | --- | --- | --- | --- |
|  | mean | SD | mean | SD | mean | SD | mean | SD | mean | SD |
| MATAM | 1.28 | 0.55 | 99.3 | 0.2 | 0.03 | 0.02 | 0.00 | 0.00 | 1252 | 116.9 |
| EMIRGE | 36.89 | 9.42 | 79.9 | 11.6 | 0.62 | 0.16 | 0.55 | 0.36 | 1436 | 15.4 |
| REAGO | 42.11 | 10.36 | 91.5 | 0.8 | 0.31 | 0.13 | 0.00 | 0.00 | 1333 | 298.9 |
| SPAdes | 21.23 | 9.05 | 73.5 | 15.9 | 0.60 | 0.49 | 0.02 | 0.04 | 966 | 47.4 |
| MEGAHIT | 23.81 | 2.85 | 80.3 | 4.9 | 0.36 | 0.18 | 0.00 | 0.00 | 962 | 87.6 |

For each of the five tools, we reported the recovered genome fraction (GF) with respect to increasing sequencing depth (Figure 2). MATAM recovered from 76% to 85% of the reference sequences for sequencing depths greater than 10x, while EMIRGE recovered less than 55% of the reference sequence, and the GF for other methods is lower than 22%. MATAM also achieved the best performance facing a low sequencing depth of 2x, reaching a GF of 33%, while GFs ranged between 5% and 10% with all other assemblers.

**Figure 2:**
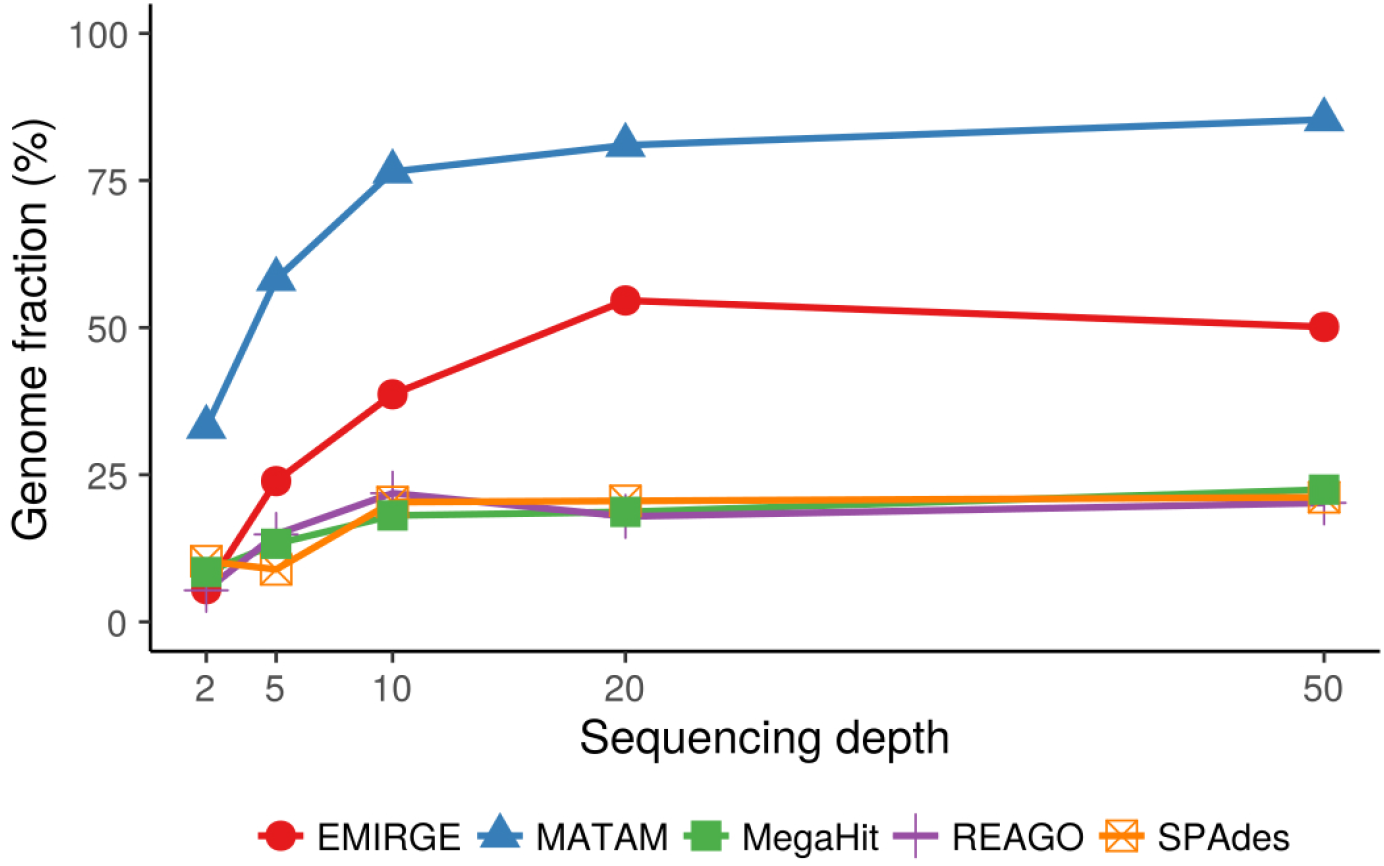
Effect of sequencing depth on the assemblies genome fractions.

### 4.2 Synthetic archaeal and bacterial community

Inching toward more realistic applications, a second dataset provides Illumina reads extracted from a synthetic microbial community composed of 16 archaeal species from 12 genera, as well as 48 bacterial species from 36 genera (accession SRR606249; [29]). As emphasized by the authors, the selected organisms cover a wide range of environmental conditions and adaptation strategies. In contrast to the previous simulated dataset (Section 4.1), the proportion of each species in the sample is not uniform, which results in individual genome average sequencing depth varying from 9x to 318x. The number of 16S rRNA paralogs per genome appears also highly diverse, ranging from 1 to 10 copies per genome. Altogether, this dataset represents a total amount of 106 distinct 16S rRNA sequences with pairwise sequence identities ranging from 59.64% to 99.93%.

The organisms were sequenced on Illumina HighSeq2000, providing 109 million 101bp paired-end reads with an average fragment size of 250bp. We quality cleaned the reads using Prinseq Lite [26], removed adapter sequences using Cutadapt [14], filtered out short reads (< 50bp), and obtained a total number of 67.6 million reads, which were analyzed with MATAM and EMIRGE. The uncleaned raw dataset was provided to REAGO, considering that the method could not handle reads with varying lengths. Finally, for SPAdes and MEGAHIT, the 16S rRNA reads were extracted from the cleaned dataset using SortMeRNA, which provided 108,560 16S rRNA reads to assemble. Cleaning and pre-processing command-lines and parameters for the synthetic community can be found in Supplementary Material (section 4.3.2)

Results are shown in Table 2. Confirming the trends observed on the simulated dataset, MATAM is able to recover the highest number of sequences together with the highest GF (83%). Most importantly, with lower ER than achieved by the other tested methods, the MATAM assembly appears highly accurate. While EMIRGE is the second best approach in terms of recovered GF, it also yields the greatest ER and Ns over all the compared tools. Moreover, a RDP classification of MATAM and EMIRGE sequences indicates that while MATAM missed one expected genus only, EMIRGE missed 4 genera out of 48.

**Table 2:** Results for the synthetic community.

|  | Chimera (%) | #contigs | TL | TAL | GF (%) | ER (%) | Ns (%) |
| --- | --- | --- | --- | --- | --- | --- | --- |
| MATAM | 3.2 | 101 | 139220 | 130654 | 83.1 | 0.05 | 0 |
| EMIRGE | 17.4 | 82 | 117138 | 102856 | 50.7 | 0.17 | 1.12 |
| REAGO | 15.5 | 59 | 90269 | 81297 | 42.8 | 0.06 | 0 |
| SPAdes | 5.5 | 59 | 70229 | 59988 | 39.9 | 0.11 | 0.05 |
| MEGAHIT | 3.0 | 61 | 77251 | 68904 | 44.3 | 0.18 | 0 |

Inspection of the MetaQuast alignments of the assemblies against the original 16S rRNAs revealed that all methods accurately assembled the genes sharing less than 90% sequence identity with their closest relatives within the sample. However, performances significantly dropped when attempting to assemble the closely related genes in the dataset. This especially concerned the paralogous 16S rRNA copies sharing around 99% sequence identity. Supplementary Table 1 (Supplementary Material, section 4.3.2) provides pairwise distances between sequences from a representative subset of four related species possessing one to three such paralogous copies. Those 16S rRNAs and their corresponding assembled candidate sequences were selected for a phylogenetic tree reconstruction. The obtained tree (Figure 3) demonstrates that MATAM correctly assembled all the different paralogs with nearly no error, while EMIRGE and REAGO only managed to recover one candidate sequence per species. Thus, EMIRGE and REAGO merged into a single candidate sequence the reads issued from distinct paralogs, resulting in erroneous assemblies with high ER and underestimated GF. Indeed, each of the sequences assembled with REAGO, as well as one EMIRGE sequence over four, appear to cluster at a slight distance from their respective targeted paralogs. Those distances simply account for the methods reconstruction errors. Consistently, in two cases, the candidates assembled by EMIRGE and REAGO were identified as chimeras by VSEARCH.

**Figure 3:**
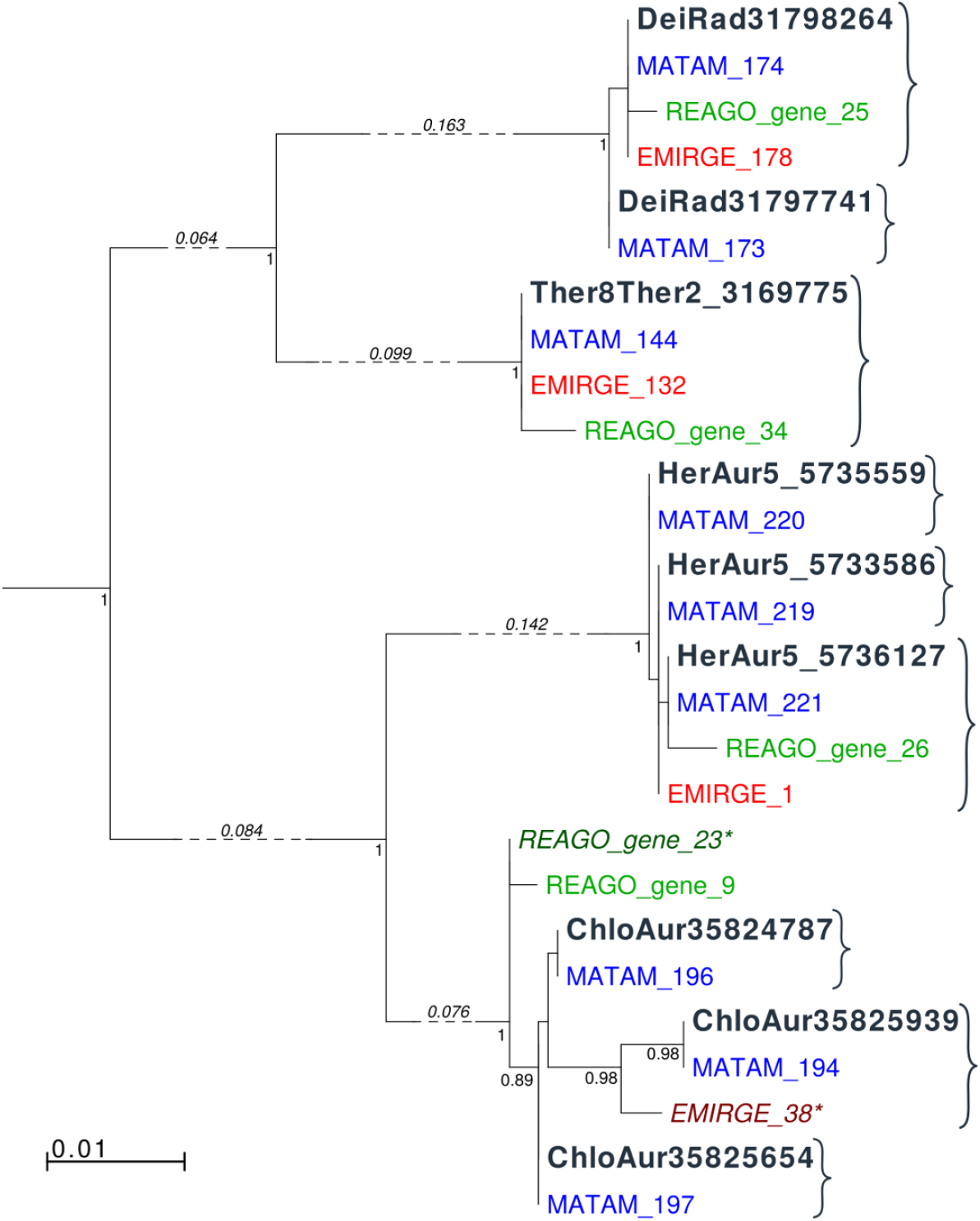
Alignment of the reference sequences with the assembled contigs shows MATAM ability to differentiate between very close sequences. MATAM, EMIRGE and REAGO contigs are shown respectively in blue, red and green. In a ideal setting, each software should produce contigs that cluster closely to each reference (black) sequence. Contigs followed by a star, and drawn in a darker color, were considered as chimeric by VSEARCH.

### 4.3 Human Microbiome Project

Finally, we used two metagenomic samples from the Human Microbiome Project (gut: SRS011405, and mouth: SRS016002, [31]) in order to validate MATAM on real metagenomic datasets sequenced from genuine environments. The reads were already quality cleaned and trimmed, and no additional filtering was performed. Hence, reads having different lengths, we were not able to run REAGO on these datasets. Results obtained with SPAdes and MEGAHIT using the following protocol appeared highly inaccurate and therefore, they are not further commented in this work. Thus, we only present the results obtained with EMIRGE and MATAM. Datasets availability, and additional details on the evaluation protocol can be found in Supplementary Material (section 4.3.3).

For these two datasets, the exact ground truth is unknown. Thus we could not perform the same validation procedure as in the two previous examples and we had to resort to alternative strategies. First, we took advantage of the availability of OTU sequences inferred through a QIIME analysis of the V1-V3 hypervariable regions for the same biological samples (available from the SRS accession numbers). We compared the assignments obtained from assemblies, calculated with RDP, with these of amplicon OTUs (Table 3). For both samples, MATAM identified more classes and genera than EMIRGE did, and most of these taxa were validated by the amplicon OTUs. Interestingly, we observed that in the two samples, three genera were recovered both by MATAM and EMIRGE, but not by the amplicon approach: *Odoribacter, Peptococcus*, and *Bergeyella.* Since some species from these genera are known to be adapted to the human gut and mouth environments, it is plausible that they were missed by the amplicon approach while being accurately recovered by MATAM and EMIRGE from the metagenomic samples.

**Table 3:** Results for the gut and mouth HMP datasets. The column *#classes* indicates the total number of taxonomic classes found with RDP from the assemblies, with the number of these classes validated with the QIIME OTUs (in parentheses). The column *#genera* gives the same information at the genus level.

|  |  | Chimera (%) | #contigs | TL | #classes | #genera |
| --- | --- | --- | --- | --- | --- | --- |
| SRS011405 | MATAM | 3.37% | 218 | 187710 | 5 (4) | 21 (17) |
|  | EMIRGE | 43.04% | 273 | 393152 | 2 (2) | 12 (8) |
| SRS016002 | MATAM | 4.92% | 353 | 320748 | 13 (13) | 31 (28) |
|  | EMIRGE | 46.01% | 282 | 394087 | 12 (12) | 25 (23) |

Moreover, we evaluated assembly quality by aligning MATAM and EMIRGE sequences against the complete Silva 128 SSU Ref NR database, using BLAST. The rationale for this experiment is that most of the species in these human gut and mouth samples are possibly already known, and therefore should be found in Silva. We observed that nearly all MATAM sequences matched with a known 16S rRNA in Silva with more than 99% identity, among which a majority matched with 100% identity (Figures 4 and 5), which suggests that MATAM sequences could possibly be assigned at the species or even the strain level. On the other hand, EMIRGE sequences provided a discordant picture. In the case of the human mouth sample, most of the EMIRGE sequences obtained a match above 97% identity, but only a slight proportion of them matched with 100% identity against a known 16S rRNA (Figure 5). The observation is even more pronounced with the human gut sample, where only 43% of the EMIRGE sequences obtained a match above 97% identity against a Silva 16S rRNA sequence (Figure 4). Thus, conversely to MATAM, EMIRGE sequences would suggest that only a slight proportion of the human gut and mouth diversity has a known isolate registered in Silva. However, considering our previous conclusions on controlled datasets, we assume that part of this diversity inferred with EMIRGE might in fact corresponds to reconstruction artifacts.

**Figure 4:**
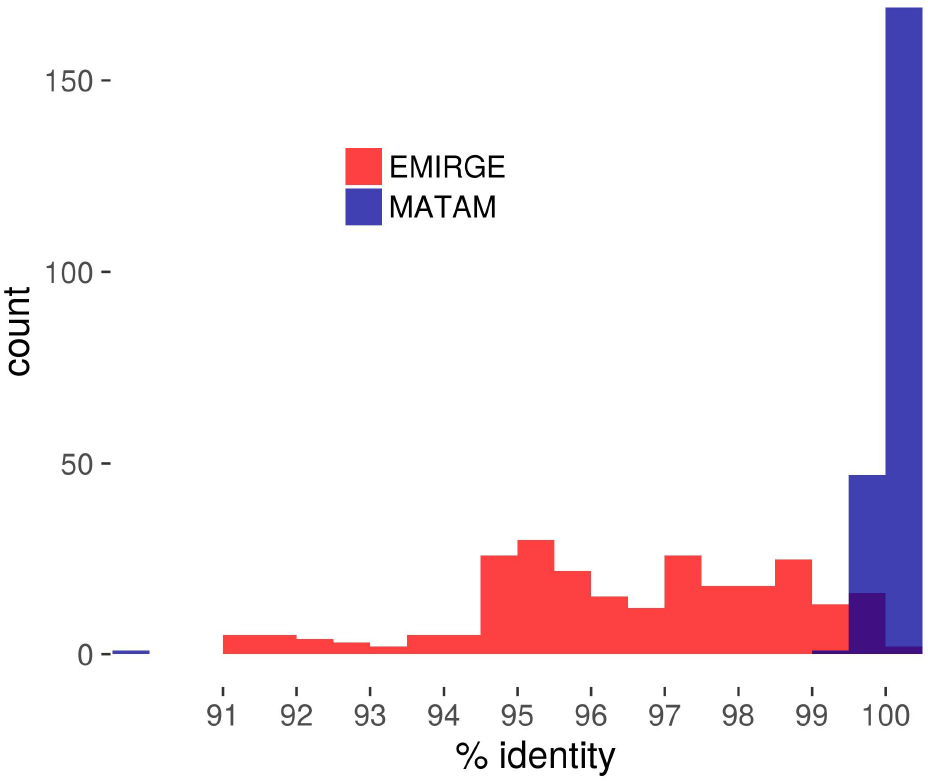
Human gut sample SRS011405. % identity distribution of best matches against Silva 128 SSU Ref NR

**Figure 5:**
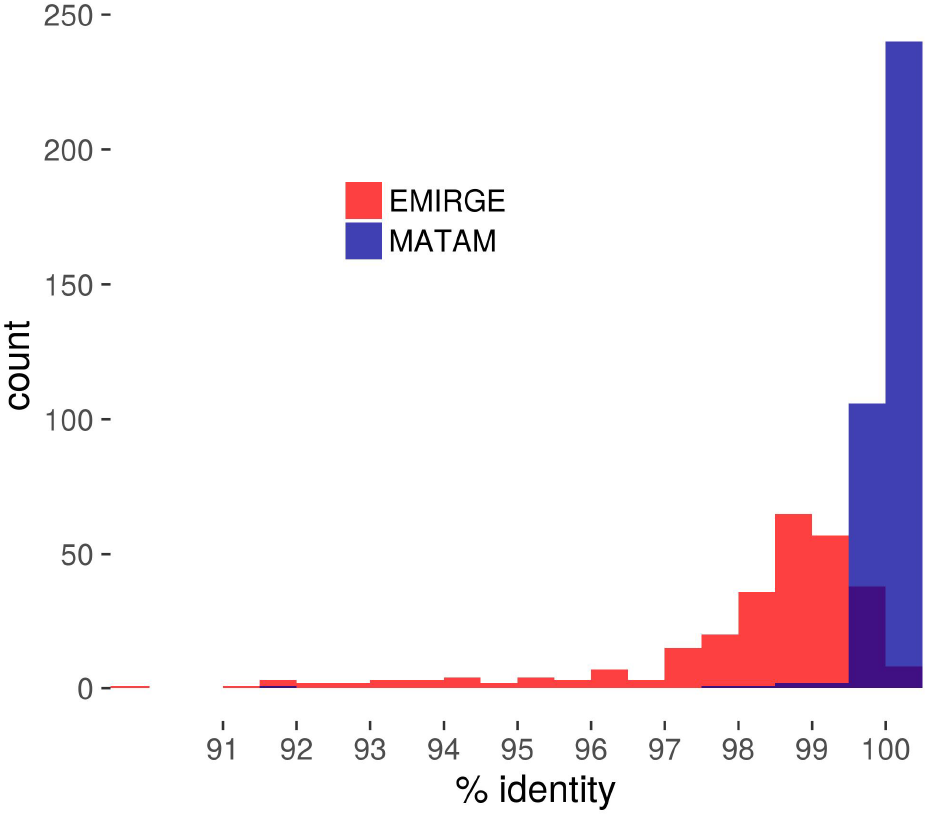
Human mouth sample SRS016002. % identity distribution of best matches against Silva 128 SSU Ref NR

## 5 Discussion

Taxonomic assignments of environmental samples is a strikingly difficult task which suffers from inherent limitations of high-throughput sequencing technologies. In this respect, we designed MATAM as an alternative to existing software helping to better understand the taxonomic structures of shotgun metagenomic samples. Our experimental results show that MATAM outperforms other available tools providing phylogenetic marker assemblies. Reconstructing full length 16S rRNAs allows to reach a higher precision of taxonomic assignments than individual read analysis or amplicon sequencing do, because the reconstructed sequences effectively contain stronger phylogenetic signal. Moreover, metagenomic shotgun sequencing is naturally immune against the primer and amplification biases attached to the amplicon sequencing technology, and therefore is more adequate to sequence unknown species.

Our approach opens up several new perspectives. Although we have focused this work on the assembly of 16S rRNA genes, MATAM was designed to deal with any marker of taxonomic interest. Indeed, there is currently an emerging trend to consider a combination of universal (single-copy) marker families, such as provided in the recently published database proGenomes [16]. Sequences from this database, or from any other customized one, could be used with MATAM to target a variety of markers, and thus provide improving taxonomic assignments. MATAM could also be used in combination with other types of sequencing data. Long read sequencing is able to produce fragments that cover large regions of the DNA molecules, up to several thousands of bases. When long reads are available, they could serve as a guide in the scaffolding step of MATAM and concomitantly, MATAM low-error contigs could be used to correct them. Finally, targeted gene capture, that allows to sequence at high depth captured DNA regions of interest from an environmental sample [6], could also prove to be an exciting application field for MATAM.

## References

[1] Anton Bankevich, Sergey Nurk, Dmitry Antipov, Alexey A. Gurevich, Mikhail Dvorkin, Alexander S. Kulikov, Valery M. Lesin, Sergey I. Nikolenko, Son Pham, Andrey D. Prjibelski, Alexey V. Pyshkin, Alexander V. Sirotkin, Nikolay Vyahhi, Glenn Tesler, Max A. Alekseyev, and Pavel A. Pevzner. SPAdes: A New Genome Assembly Algorithm and Its Applications to Single-Cell Sequencing. Journal of Computational Biology, 19(5):455–477, May 2012.

[2] James R. Cole, Qiong Wang, Jordan A. Fish, Benli Chai, Donna M. McGarrell, Yanni Sun, C. Titus Brown, Andrea Porras-Alfaro, Cheryl R. Kuske, and James M. Tiedje. Ribosomal Database Project: data and tools for high throughput rRNA analysis. Nucleic Acids Research, 42(D1):D633–D642, January 2014.

[3] T. Z. DeSantis, P. Hugenholtz, N. Larsen, M. Rojas, E. L. Brodie, K. Keller, T. Huber, D. Dalevi, P. Hu, and G. L. Andersen. Greengenes, a Chimera-Checked 16s rRNA Gene Database and Workbench Compatible with ARB. Applied and Environmental Microbiology, 72(7):5069–5072, January 2006.

[4] Andreas Döring, David Weese, Tobias Rausch, and Knut Reinert. Seqan an efficient, generic c++ library for sequence analysis. BMC Bioinformatics, 9(1):1–9, 2008.

[5] Robert C. Edgar, Brian J. Haas, Jose C. Clemente, Christopher Quince, and Rob Knight. UCHIME improves sensitivity and speed of chimera detection. Bioinformatics, 27(16):2194–2200, August 2011.

[6] Cyrielle Gasc, Eric Peyretaillade, and Pierre Peyret. Sequence capture by hybridization to explore modern and ancient genomic diversity in model and nonmodel organisms. Nucleic Acids Research, 44(10):4504–4518, June 2016.

[7] Weichun Huang, Leping Li, Jason R. Myers, and Gabor T. Marth. ART: a next-generation sequencing read simulator. Bioinformatics, 28(4):593–594, February 2012.

[8] Evguenia Kopylova, Jose A. Navas-Molina, Céline Mercier, Zhenjiang Zech Xu, Frédéric Mahé, Yan He, Hong-Wei Zhou, Torbjørn Rognes, J. Gregory Caporaso, and Rob Knight. Open-source sequence clustering methods improve the state of the art. mSystems, 1(1), 2016.

[9] Evguenia Kopylova, Laurent Noé, Pierre Pericard, Mikael Salson, and Hélène Touzet. Sortmerna 2: ribosomal rna classification for taxonomic assignation. In Workshop on Recent Computational Advances in Metagenomics, ECCB 2014, 2014.

[10] Evguenia Kopylova, Laurent Noé, and Hélène Touzet. Sortmerna: fast and accurate filtering of ribosomal rnas in metatranscriptomic data. Bioinformatics, 28(24):3211–3217, 2012.

[11] Dinghua Li, Chi-Man Liu, Ruibang Luo, Kunihiko Sadakane, and Tak-Wah Lam. MEGAHIT: an ultra-fast single-node solution for large and complex metagenomics assembly via succinct de Bruijn graph. Bioinformatics, 31(10):1674–1676, May 2015.

[12] Bo Liu, Theodore Gibbons, Mohammad Ghodsi, Todd Treangen, and Mihai Pop. Accurate and fast estimation of taxonomic profiles from metagenomic shotgun sequences. BMC Genomics, 12(Suppl 2):S4, July 2011.

[13] Kenneth J. Locey and Jay T. Lennon. Scaling laws predict global microbial diversity. Proceedings of the National Academy of Sciences, 113(21):5970–5975, May 2016.

[14] Marcel Martin. Cutadapt removes adapter sequences from high-throughput sequencing reads. EMBnet. journal, 17(1):pp. 10–12, May 2011.

[15] Konstantinos Mavromatis, Natalia Ivanova, Kerrie Barry, Harris Shapiro, Eugene Goltsman, Alice C. McHardy, Isidore Rigoutsos, Asaf Salamov, Frank Korzeniewski, Miriam Land, Alla Lapidus, Igor Grigoriev, Paul Richardson, Philip Hugenholtz, and Nikos C. Kyrpides. Use of simulated data sets to evaluate the fidelity of metagenomic processing methods. Nature Methods, 4(6):495–500, June 2007.

[16] Daniel R. Mende, Ivica Letunic, Jaime Huerta-Cepas, Simone S. Li, Kristoffer Forslund, Shinichi Sunagawa, and Peer Bork. proGenomes: a resource for consistent functional and taxonomic annotations of prokaryotic genomes. Nucleic Acids Research, 45(D1):D529–D534, January 2017.

[17] Celine Mercier, Frederic Boyer, Aurelie Bonin, and Coissac Eric. Sumatra and sumaclust: fast and exact comparison and clustering of sequences, 2013.

[18] Alla Mikheenko, Vladislav Saveliev, and Alexey Gurevich. MetaQUAST: evaluation of metagenome assemblies. Bioinformatics, 32(7):1088–1090, April 2016.

[19] Christopher S. Miller, Brett J. Baker, Brian C. Thomas, Steven W. Singer, and Jillian F. Banfield. Emirge: reconstruction of full-length ribosomal genes from microbial community short read sequencing data. Genome Biology, 12(5):R44, 2011.

[20] Eric P. Nawrocki, Diana L. Kolbe, and Sean R. Eddy. Infernal 1.0: inference of RNA alignments. Bioinformatics, 25(10):1335–1337, May 2009.

[21] Sergey Nurk, Dmitry Meleshko, Anton Korobeynikov, and Pavel Pevzner. metaSPAdes: a new versatile de novo metagenomics assembler. arXiv:1604.03071 [q-bio], April 2016. arXiv: 1604.03071.

[22] Miguel Pignatelli and Andrés Moya. Evaluating the Fidelity of De Novo Short Read Metagenomic Assembly Using Simulated Data. PLOS ONE, 6(5):e19984, May 2011.

[23] Rachel Poretsky, Luis M. Rodriguez-R, Chengwei Luo, Despina Tsementzi, and Konstantinos T. Konstantinidis. Strengths and Limitations of 16s rRNA Gene Amplicon Sequencing in Revealing Temporal Microbial Community Dynamics. PLoS ONE, 9(4):e93827, April 2014.

[24] Christian Quast, Elmar Pruesse, Pelin Yilmaz, Jan Gerken, Timmy Schweer, Pablo Yarza, Jörg Peplies, and Frank Oliver Glöckner. The SILVA ribosomal RNA gene database project: improved data processing and web-based tools. Nucleic Acids Research, 41(D1):D590–D596, January 2013.

[25] Torbjørn Rognes, Tomáš Flouri, Ben Nichols, Christopher Quince, and Frédéric Mahé. VSEARCH: a versatile open source tool for metagenomics. PeerJ, 4:e2584, October 2016.

[26] Robert Schmieder and Robert Edwards. Quality control and preprocessing of metagenomic datasets. Bioinformatics, 27(6):863–864, March 2011.

[27] Alexander Sczyrba, Peter Hofmann, Peter Belmann, David Koslicki, Stefan Janssen, Johannes Droege, Ivan Gregor, Stephan Majda, Jessika Fiedler, Eik Dahms, Andreas Bremges, Adrian Fritz, Ruben Garrido-Oter, Tue Sparholt Jorgensen, Nicole Shapiro, Philip D. Blood, Alexey Gurevich, Yang Bai, Dmitrij Turaev, Matthew Z. DeMaere, Rayan Chikhi, Niranjan Nagarajan, Christopher Quince, Lars Hestbjerg Hansen, Soren J. Sorensen, Burton K. H. Chia, Bertrand Denis, Jeff L. Froula, Zhong Wang, Robert Egan, Dongwan Don Kang, Jeffrey J. Cook, Charles Deltel, Michael Beckstette, Claire Lemaitre, Pierre Peterlongo, Guillaume Rizk, Dominique Lavenier, Yu-Wei Wu, Steven W. Singer, Chirag Jain, Marc Strous, Heiner Klingenberg, Peter Meinicke, Michael Barton, Thomas Lingner, Hsin-Hung Lin, Yu-Chieh Liao, Genivaldo Gueiros Z. Silva, Daniel A. Cuevas, Robert A. Edwards, Surya Saha, Vitor C. Piro, Bernhard Y. Renard, Mihai Pop, Hans-Peter Klenk, Markus Goeker, Nikos Kyrpides, Tanja Woyke, Julia A. Vorholt, Paul Schulze-Lefert, Edward M. Rubin, Aaron E. Darling, Thomas Rattei, and Alice C. McHardy. Critical Assessment of Metagenome Interpretation - a benchmark of computational metagenomics software. bioRxiv, p. 099127, January 2017.

[28] Nicola Segata, Levi Waldron, Annalisa Ballarini, Vagheesh Narasimhan, Olivier Jousson, and Curtis Huttenhower. Metagenomic microbial community profiling using unique clade-specific marker genes. Nature Methods, 9(8):811–814, 2012.

[29] Migun Shakya, Christopher Quince, James H. Campbell, Zamin K. Yang, Christopher W. Schadt, and Mircea Podar. Comparative metagenomic and rRNA microbial diversity characterization using Archaeal and Bacterial synthetic communities. Environmental microbiology, 15(6):1882–1899, June 2013.

[30] Jared T. Simpson and Richard Durbin. Efficient de novo assembly of large genomes using compressed data structures. Genome Research, 22(3):549–556, 2012.

[31] The Human Microbiome Project Consortium. Structure, function and diversity of the healthy human microbiome. Nature, 486(7402):207–214, June 2012.

[32] Qiong Wang, George M. Garrity, James M. Tiedje, and James R. Cole. Naïve Bayesian Classifier for Rapid Assignment of rRNA Sequences into the New Bacterial Taxonomy. Applied and Environmental Microbiology, 73(16):5261–5267, August 2007.

[33] Cheng Yuan, Jikai Lei, James Cole, and Yanni Sun. Reconstructing 16s rRNA genes in metagenomic data. Bioinformatics, 31(12):i35–i43, June 2015.

